# Cleavage of MEP-1 by DPF-3 Reveals Novel Substrate Specificity and Its Impact on Reproductive Fitness

**DOI:** 10.1101/2025.09.26.678732

**Authors:** Ilkin Aygün, Afzal Amanullah, Jan Seebacher, Daniel Hess, Charlotte Soneson, Helge Großhans, Rajani Kanth Gudipati

## Abstract

Proteases are enzymes that catalyse the hydrolysis of peptide bonds in proteins for their functional, modification or degradation. Members of the Dipeptidyl Peptidase IV (DPPIV) family are exopeptidases that cleave dipeptides off the N-termini of their substrate peptides, typically after proline or alanine. Recently, we showed that human DPP4 and *Caenorhabditis elegans* DPF-3 have a larger target repertoire *in vitro*, permitting cleavage after additional amino acids. Here, we use TAILS (Terminal Amine Isotopic Labelling of Substrates) to identify DPF-3 targets *in vivo* and observe cleavage of MEP-1 after threonine, confirming a broader substrate specificity of DPF-3 also *in vivo*. Demonstrating physiological relevance, we show that rendering MEP-1 resistant to cleavage disrupts its stability, leading to developmental abnormalities such as defective gonadal migration and reproductive issues. Collectively, our findings highlight a previously unappreciated complexity in the substrate specificity of DPPIV family proteases and suggest that their physiological roles may extend beyond what is currently known.

**IMPORTANT:**

- Manuscripts submitted to Review Commons are peer reviewed in a journal-agnostic way.
- Upon transfer of the peer reviewed preprint to a journal, the referee reports will be available in full to the handling editor.
- The identity of the referees will NOT be communicated to the authors unless the reviewers choose to sign their report.
- The identity of the referee will be confidentially disclosed to any affiliate journals to which the manuscript is transferred.

**GUIDELINES:**

- For reviewers: https://www.reviewcommons.org/reviewers
- For authors: https://www.reviewcommons.org/authors

**CONTACT:** The Review Commons office can be contacted directly at

## Introduction

Proteases are enzymes that catalyze the hydrolysis of the peptide bonds in proteins, generating smaller peptides or individual amino acids. The proteases are expressed in all kingdoms of life and were initially believed to be restricted to the purpose of digestion and recycling of the amino acids, but later research has shown their importance in several cellular processes. They play a pivotal role in both physiological and pathological processes, making them valuable targets for drug development (Lopez-Otin and Bond 2008).

Proteases are classified into several groups based on their catalytic residue (Lopez-Otin and Bond 2008). The Dipeptidyl peptidase IV (DPPIV) family proteins are serine class proteases belonging to the S9b protease subfamily, with a conserved catalytic triad comprised of Serine, Aspartic acid and Histidine (Hama et al. 1982; Hopsu-Havu and Glenner 1966). In mammals, this family is comprised of four enzymatically active members, DPP4, DPP8, DPP9 and FAP (Fibroblast Activation Protein) and two enzymatically inactive members, DPP6 and DPP10, which lack the catalytic serine (Yu et al. 2010). Identification of the substrates and elucidating the molecular mechanism/s of action of proteases aid in the development of medication for human benefit (Drag and Salvesen 2010). The DPPIV family proteases are of clinical significance, given their role in glucose metabolism. It was identified that GLP-1, an incretin hormone that stimulates insulin secretion (Komatsu et al. 1989) is a substrate of DPP4 (Mentlein, Gallwitz, and Schmidt 1993). Over several decades, specific inhibitors of DPP4 have been developed and are widely used to treat type II diabetes (Deacon 2019; Demuth, McIntosh, and Pederson 2005). Inhibition of DPP8/9 leads to pyroptosis (Okondo et al. 2018) and inhibitors of DPP8/9 could potentially be used to treat acute myeloid leukemia (Johnson et al. 2018). In spite of research over several decades on DPPIV family proteases, only the inhibitors of DPP4 have successfully been developed for human use, largely aided by prior identification of its substrates. Lack of knowledge regarding the substrate repertoire/s of DPPIV family proteases remains one of the major bottlenecks to deeply understand their mechanism/s of action and further development of their inhibitors (Mulvihill and Drucker 2014; Tagore et al. 2009; Tinoco, Tagore, and Saghatelian 2010; Wilson et al. 2016).

The genome of *Caenorhabditis elegans* encodes seven *dpf* genes (*dpf-1* to *dpf-7*), among which the *dpf-1* through *dpf-6* contain the catalytic triad, suggesting that these members of the DPPIV family are likely active proteases (Gudipati et al. 2021). We have earlier shown that the *dpf-3*, the *C. elegans* ortholog of mammalian DPP8/9 is required for reproduction. Specifically, we showed that the RNA binding proteins WAGO-1 and WAGO-3 are substrates of DPF-3. Loss of DPF-3 catalytic activity leads to depletion of WAGO-3 protein levels, while WAGO-1 becomes loaded with non-canonical short interfering RNAs (siRNAs). This misregulation results in the de-silencing of transposable elements, increased DNA damage and fully penetrant male sterility because of defective spermatogenesis (Gudipati et al. 2021). Recently, we have further shown that the DPF-3 physically and genetically interacts with ALG-1, an Argonaute protein that is central for the microRNA mediated gene expression regulation. The loss of *dpf-3* suppresses the *alg-1* loss of function by increasing the levels of ALG-2, a paralog of ALG-1 (Harvey et al. 2025), through a mechanism that is independent of DPF-3’s catalytic activity.

DPP4, DPP8, and DPP9 are presumed to be strictly N-terminal dipeptidyl peptidases (Heymann and Mentlein 1978; Mentlein, Gallwitz, and Schmidt 1993; Yu et al. 2009). The DPPIV family proteases are also referred to as prolyl peptidases because of their preference for proline (Pro) at the P1 position in their substrates (Keane et al. 2011) (The second amino acid position from the N-terminus is referred to as the P1 position and the cleavage occurs at the C-terminus of the P1 position). They also tolerate alanine (Ala) and liberates Xaa-Pro or Xaa-Ala from their substrates. Although it has been proposed that the DPPIV proteases accept smaller amino acids such as glycine (Gly), serine (Ser), valine (Val) at the P1 position (Deacon 2019), physiological substrates containing these non-canonical amino acids at the P1 position are yet to be identified. Using an *in vitro* approach named, qPISA, we have recently shown that DPP4 and DPF-3 can also accept Ser, threonine (Thr) and Gly at P1 position (Gudipati et al. 2024). However, no physiological substrates with residues other than Pro or Ala at the P1 position have been identified for any of the DPPIV family proteases in any model organism. Importantly, the functional consequences of cleaving substrates at non-canonical amino acids, or of failing to do so, remain unexplored.

Here, using an unbiased proteome wide approach, we report the identification of the substrates of DPF-3 in developing nematodes using TAILS (Terminal Amine Isotopic Labelling of Substrates), to enrich the N-termini of the proteome (Kleifeld et al. 2011). Our approach identified several canonical substrates (having Pro or Ala at the P1 position) as well as a non-canonical substrate, MEP-1, containing Thr at the P1 position that we confirmed through an independent assay. Mutating the DPF-3 recognition site in MEP-1 rendering it refractive to catalytic activity leads to a decrease in its steady state level culminating in gonadal migration defects and subsequent reproductive impairment. The new results reported here broaden our understanding of DPF-3 and shed further light on its role in development and reproduction.

## Results

### Identification of DPF-3 *in vivo* substrates

Through a targeted approach, we previously determined that WAGO-1 and WAGO-3 are key substrates of DPF-3 in controlling genome integrity (Gudipati et al. 2021). We wondered if there are additional substrates of DPF-3, possibly linked to other functions, and aimed to identify them through an unbiased approach. Intriguingly, our recent *in vitro* work suggested that DPF-3 can accept non-canonical amino acids such as Thr or Ser at the P1 position in their substrates (Gudipati et al. 2024). This agrees with similar observations for mammalian DPPIV family proteases (Deacon 2019), but physiological substrates of this type remain to be identified. To determine the substrates and to discover potential substrates having non-canonical amino acids at the P1 position *in vivo*, we chose a mass spectrometry based method, named, TAILS that enriches the N-termini of the proteins (Kleifeld et al. 2011) (Fig.1A).

**Fig. 1.**
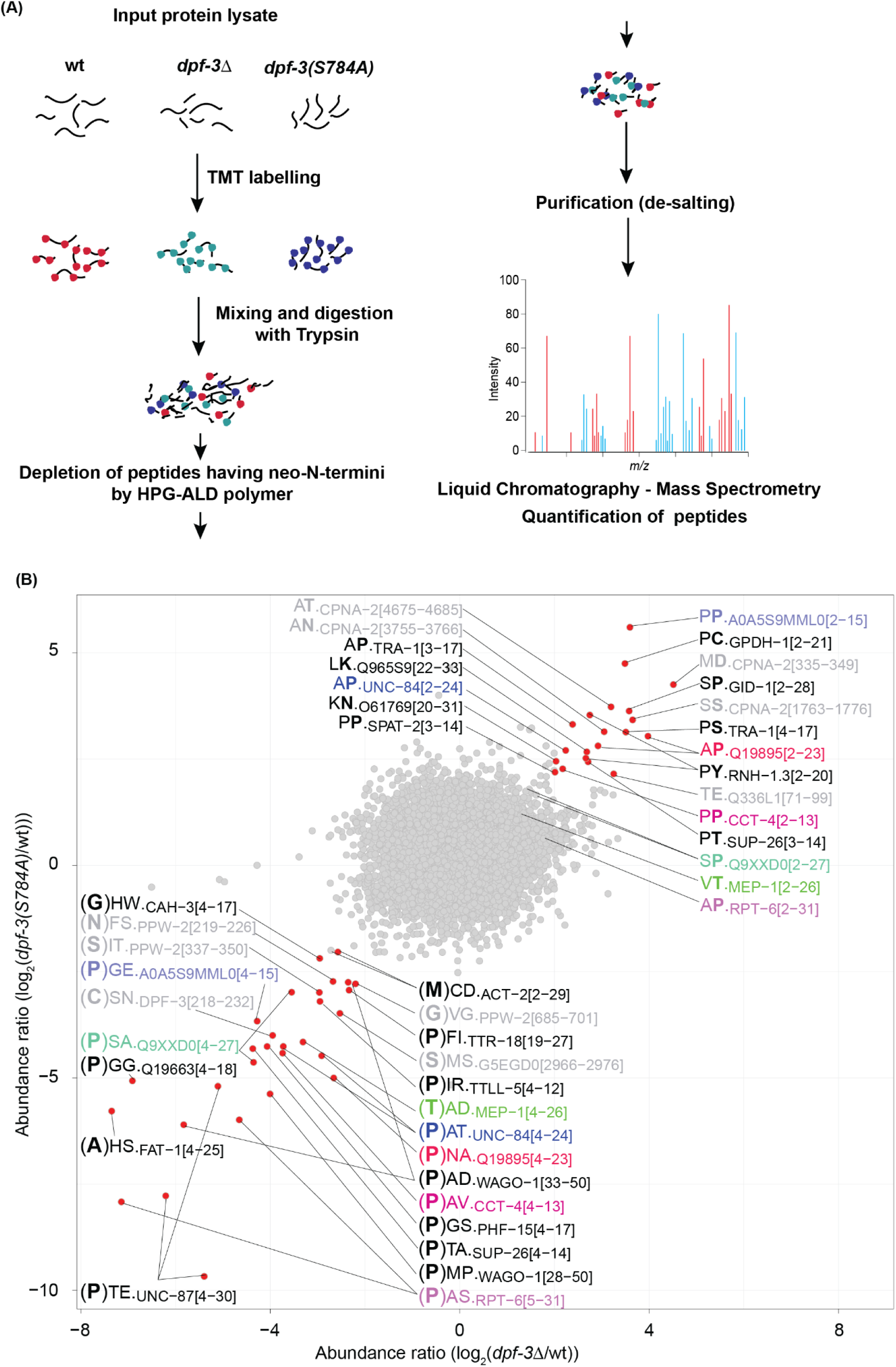
Identification of *in vivo* substrates of DPF-3 A. Cartoon representing the TAILS protocol. B. Scatter plot showing the log_2_-transformed abundance ratio of the detected peptides in indicated mutant relative to wild-type control animals after enrichment of N-termini by TAILS. Each dot represents a peptide. Peptides exhibiting an absolute log_2_-fold change of >2 in each sample comparison are colored in red and labeled with the first two amino acids from the N-terminus of the detected peptide, along with the name and position (in square brackets, relative to expected N-terminus) of the protein from which it derives. For putative products (lower left quadrant), the residue preceding the detected peptide is indicated in bold and in brackets. For putative substrates (upper right quadrant), the second amino acid (corresponding to the P1 position) is indicated in bold. Peptides originating from >50^th^ position in their cognate proteins are colored gray. Peptide pairs (peptides from same cognate protein that differ at their N-terminus depending on the genetic background, with at least one exhibiting an absolute log_2_ fold change of >2 and the matching two peptides being separated by a Manhattan distance of 4) are indicated in identical color. For instance, a peptide originating from the CCT-4 N-terminus (pink; position 2 relative to initiator Met) is increased in abundance in *dpf-3* mutant animals, whereas its matching partner (starting at position 4) is decreased in abundance. Peptides lacking a corresponding “partner” are labeled in black. Multiple peptides with the same annotation represent differences in the charge/modification status of that peptide.

We prepared total worm extracts from synchronized young adult stage animals of three different genotypes: wild-type, *dpf-3(xe68)* (almost the entire gene coding for *dpf-3* is deleted), hereafter referred to as *dpf-3Δ*, and *dpf-3(xe71[S784A])*, which exhibits severely reduced or entirely absent DPF-3 protease activity (Gudipati et al. 2021) due to the substitution of the catalytic Ser 784 to Ala. In this strain, henceforth referred to as *dpf-3(S784A)*, DPF-3 is additionally FLAG-HA epitope-tagged at its C-terminus.

We quantified ∼20,000 peptides in each of the wild-type and mutant samples (Suppl. Table 1) representing about 5,000 proteins. Consistent with the enrichment of N-termini by TAILS, the largest bin (∼1,300 per experiment) of peptides aligned within the first 5 amino acids window of the annotated protein sequence. The next two largest bins were starting at amino acids position 16-20 and 21-25, respectively, likely reflecting the removal of a signal peptide from the N-termini of the nascent proteins in this bin (Suppl. Fig. 1).

When analyzing peptide changes in each mutant relative to the corresponding wild-type sample, we anticipated observing a depletion of products (semi-tryptic peptides shortened by 2 amino acids compared to the parental tryptic peptide) in the *dpf-3* mutant strains relative to wild-type. Conversely, in wild-type lysates, substrates (semi-tryptic peptides representing the N-terminus of the cognate protein) were expected to show depletion alongside an increase in products (shortened by two amino acids). Ideally, consistent substrate-product pairs should be observed; however, several technical and biological factors can disrupt this pattern. Technical factors include limitations such as the inherent detection bias of mass spectrometry, which may prevent the identification of specific peptides.

Biological factors include variations in the stability and degradation rates of substrates and products, which can influence their relative abundance and detection. Efficient substrate turnover can result in rapid substrate depletion and an abundance of product. On the other hand, if a substrate is poorly converted to its product, there may be little to no observable depletion of the substrate. However, even a small accumulation of the product can show a large fold change (from negligible levels to measurable amounts), leading to a more pronounced change in the product relative to the substrate. Such patterns typically arise when enzymatic activity or processing steps are inefficient or slow, creating a bottleneck in the reaction.

When plotting log_2_-fold changes comparing wild type to the *dpf-3* loss-of-function strains, we found two sets of peptides that changed consistently either up or down (Fig. 1B). Many of the peptides in the lower left quadrant (enriched in wild-type relative to *dpf-3* mutant animals), were two amino acids shorter than the expected protein N-terminus, consistent with their being products of DPF-3 processing. The N-terminus is typically defined by the annotated initiator methionine unless the second amino acid is Gly, Ala, Ser, Thr, Cys, Pro, or Val. These residues promote the removal of the initiator Met, designating the second amino acid as the N-terminus (Wingfield 2017; Xiao et al. 2010). Similarly, in cases where an annotated signal sequence is present, it will be cleaved from the nascent peptide, thereby redefining the N-terminus (Martoglio and Dobberstein 1998). Conversely, peptides present in the upper right quadrant and thus enriched in *dpf-3* mutant animals typically started at the expected N-terminus, consistent with their being substrates of DPF-3 mediated processing.

We have observed several instances of matching pairs of substrates and products (Fig. 1B, Suppl. Table 2). In addition to identifying novel substrates, we also detected WAGO-1, a previously known substrate of DPF-3 (Gudipati et al. 2021). Interestingly, the peptides corresponding to WAGO-1 originate not from the N-terminus but rather from downstream positions (starting at the residues 28 and 33), consistent with sequential processing of WAGO-1 by DPF-3 (Gudipati et al. 2021).

We independently verified several of the potential substrates in an *in vitro* assay using synthetic peptides representing the N-terminus of the substrate protein and recombinant DPF-3. As shown in Fig. 2, product generation over time could be observed when the peptides were incubated with wild-type DPF-3 but not DPF-3(S784A) mutant protein. These experiments led to the identification of substrates of DPF-3 and further validation using an independent *in vitro* approach.

**Fig. 2.**
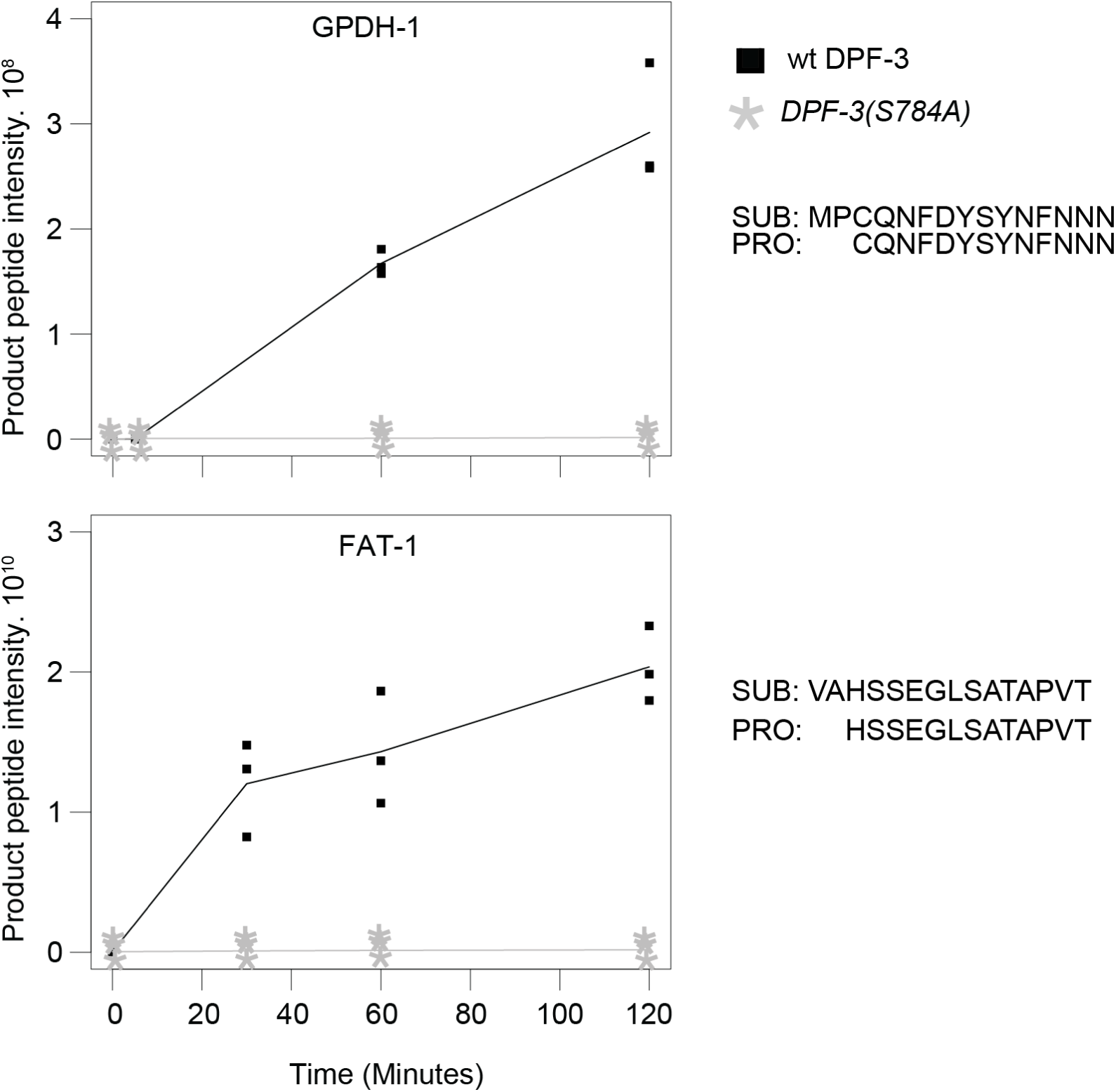
Confirmation of potential DPF-3 substrates by *in vitro* digestion Product peptide intensity of potential substrates (y-axis) representing the N-termini of GPDH-1 and FAT-1 (without the initiator methionine) is plotted over reaction time (x-axis). Data points of three technical replicates are shown; a line indicates the mean values. Since no product was detected when the DPF-3(S784A) mutant protein was used, all relevant replicate points are on top of each other. The sequences of the substrate (SUB) and the product (PRO) are shown.

### MEP-1, containing Thr at the P1 position is a substrate of DPF-3

Consistent with the *in vitro* preferences of DPF-3 revealed by qPISA (Gudipati et al. 2024), the *in vivo* processing events primarily occurred after Pro in the P1 position (Fig. 1B).

However, we also identified MEP-1, with Thr at the P1 position, as a potential substrate of DPF-3. Specifically, we detected an N-terminal peptide, MEP-1_4-26,_ whose abundance was significantly reduced in *dpf-3* mutant lysates, constituting a potential product. Conversely, the levels of the cognate MEP-1_2-26_ substrate peptide were modestly increased in the mutant lysates. This substrate contains P2(Val) and P1(Thr) as positive determinants of cleavage, while P1’(Ala) is mildly negative according to the qPISA model (Gudipati et al. 2024). The significantly larger changes observed for the product compared to the substrate align with the earlier explanation of inefficient substrate processing. This observation is consistent with predictions based on the *in vitro* qPISA data.

Since physiological substrates having Thr at P1 have not previously been reported for DPF-3 or any of the mammalian DPPIV family proteases, we sought to further validate MEP-1 as a legitimate substrate of DPF-3. Incubation of a synthetic MEP-1_2-13_ peptide with recombinant DPF-3 yielded product accumulation, whereas incubation of a *T3E*-mutated MEP-1_2-13_ control peptide did not (Fig. 3A, B). To confirm that DPF-3 processes MEP-1 *in vivo*, we used CRISPR/Cas9 mediated genome editing to generate *mep-1(xe312[3xflag::linker::mep-1])*, referred to as FLAG::MEP-1 here on (Fig. 3C). We immunopurified the MEP-1 from total lysate of synchronized young adult stage wild-type and *dpf-3Δ* animals and mapped the protein N-terminus through mass spectrometry. While internal peptide intensities, used as a reference, did not differ between wild-type and *dpf-3Δ* animals, the N-terminal substrate peptide, starting at V2, was enriched in *dpf-3Δ* mutant relative to wild-type animals. Conversely, the product peptide, starting at A4, was abundant in wild-type but barely detected in *dpf-3Δ* mutant animals (Fig. 3D). These results confirm MEP-1 as a *bona fide* substrate of DPF-3 and demonstrate DPF-3’s capacity to process proteins with Thr at the P1 position *in vivo*.

**Fig. 3.**
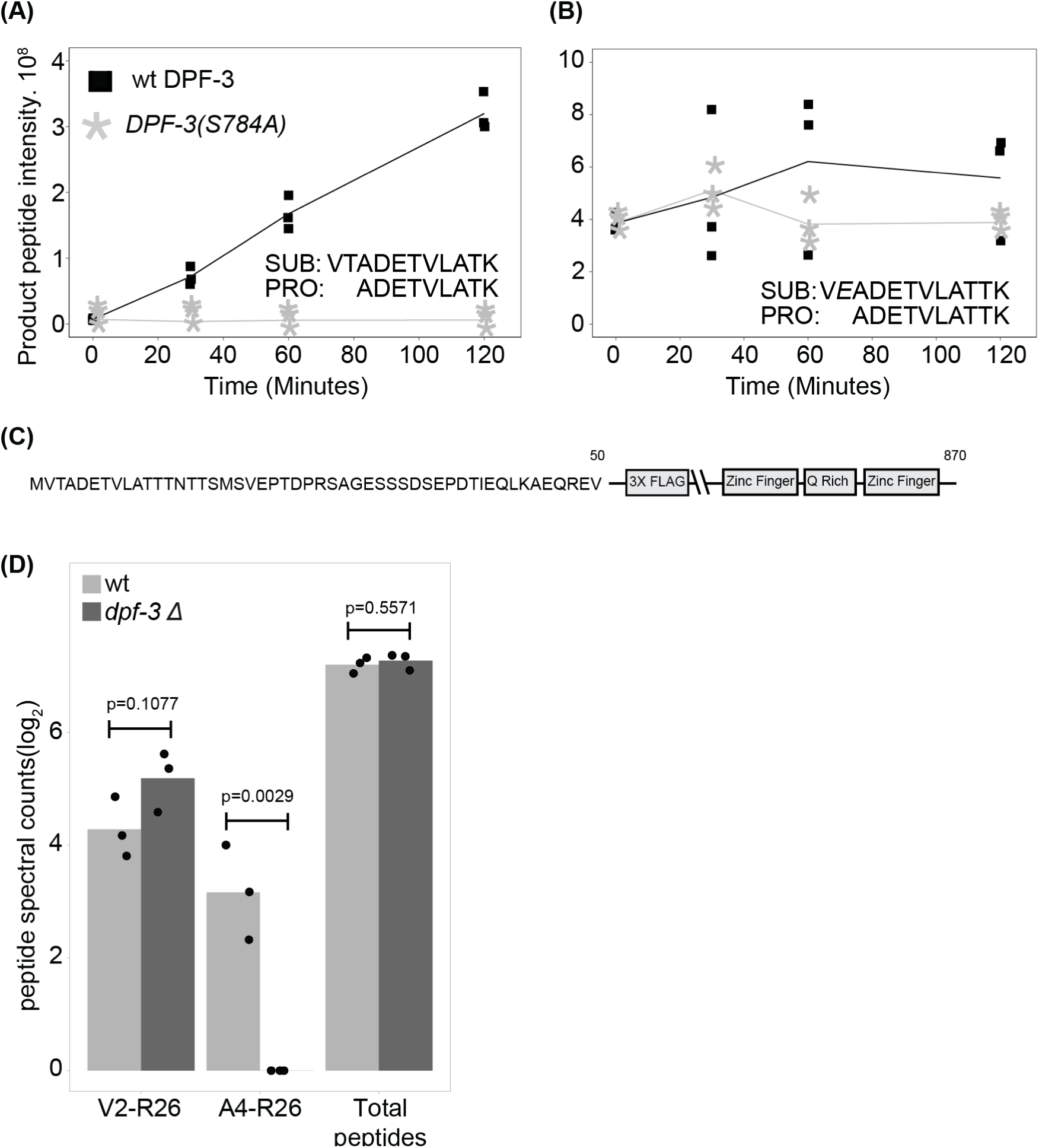
MEP-1 is a substrate of DPF-3 **A.** Product peptide intensity of a substrate (y-axis) representing the N-terminus (without the initiator methionine) of MEP-1, plotted over reaction time (x-axis). Data points of three technical replicates are shown; a line indicates the mean values. Since no product was seen when DPF-3(S784A) mutant protein was used, all the relevant replicate points are on top of each other. The sequences of the substrate (SUB) and the product (PRO) are shown. **B.** Same as panel **A**, except mutant MEP-1 (Threonine at P1 position replaced by Glutamine) peptide was used. **C.** Schematic representation of the MEP-1 protein, highlighting the transgenic *3x flag* epitope, Zinc finger domains, and the Glutamine-rich region (not drawn to scale). Numbers above the schematic indicate the corresponding amino acid positions within the native protein sequence. **D.** Bar plot of indicated peptides showing log_2_-transformed peptide spectral counts from three biological replicates of wild-type or *dpf-3Δ* mutant animals. Each dot represents a biological replicate, the bar is drawn using the average value.

### Mutation of DPF-3 recognition site in MEP-1 alters its steady state level

MEP-1 is a Kruppel-type zinc finger protein essential for preventing the development of germline cells in the somatic tissue, thereby maintaining the distinction between somatic and germline lineages (Unhavaithaya et al. 2002). Upon confirming that MEP-1 is a substrate of DPF-3 and undergoes processing *in vivo*, we sought to investigate the functional significance of this processing by generating *mep-1(xe311aga19[3xflag::linker::mep-1(T3E)])* mutant animals. To avoid the epitope interfering with the processing of the host protein, the sequence coding *3xflag::linker* is inserted away from the N terminus (between the 52^nd^ and 53^rd^ amino acids). In this allele, threonine at position 3 is replaced with glutamic acid, rendering the MEP-1 protein resistant to cleavage by DPF-3; these animals are hereafter referred to as *flag::mep-1(T3E)*. We first assessed its steady state levels by western blot. As can be seen in Fig. 4A, loss of *dpf-3* leads to a moderate decrease in MEP-1 steady state level. Interestingly, the *flag::mep-1(T3E)* mutant animals exhibited a severe depletion of MEP-1 protein. This suggests that its processing by DPF-3 removes a potential degron signal at its N-terminus, thereby stabilizing the protein. Consistent with this interpretation, quantification of *mep-1* mRNA levels by reverse transcription-qPCR (RT-qPCR) revealed no significant changes at the transcript level (Suppl. Fig. 2A).

**Fig. 4.**
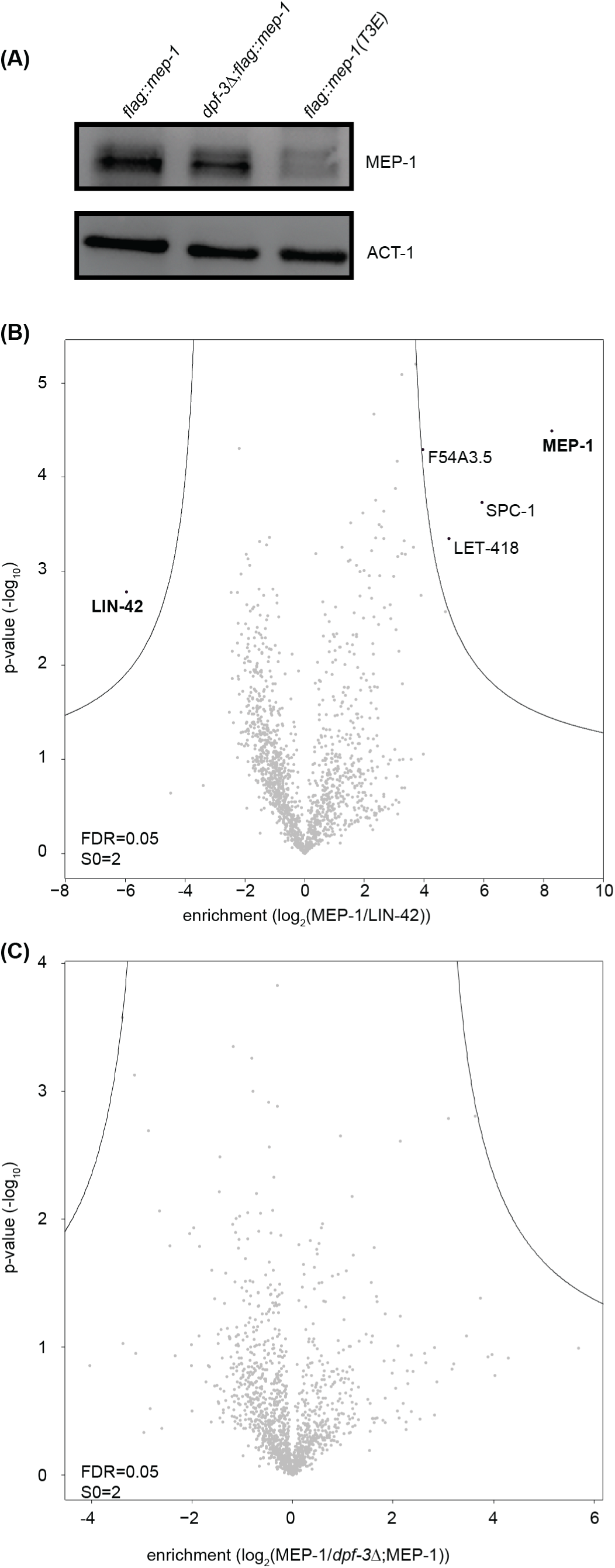
DPF-3 processing stabilizes MEP-1 without altering complex composition. **A.** Western blot showing reduced MEP-1 levels in *dpf-3Δ* animals and marked depletion of cleavage-resistant *MEP-1(T3E)* mutant protein. **B.** IP-MS of MEP-1. The x-axis indicates the fold enrichment of the bait protein (MEP-1) and its interacting partners over control (LIN-42). Each dot represents a protein, the non-significant and the significant proteins are colored gray and black, respectively. False Discovery Rate (FDR) of 0.05 was used. The Y-axis of the volcano plot indicates the p-values that were calculated by two-tailed Student t-test. **C.** Exactly as shown for B, except the comparison is between MEP-1 immunoprecipitated from the lysate of wild type or *dpf-3Δ* mutant animals.

It has been shown that MEP-1 interacts with LET-418, the *C. elegans* homolog of Mi-2/CHD3, which belongs to a family of evolutionarily conserved chromodomain proteins involved in chromatin remodeling and transcriptional repression (Unhavaithaya et al. 2002). MEP-1 also associates with HDA-1 (Unhavaithaya et al. 2002), the *C. elegans* ortholog of HDAC-1, a conserved histone deacetylase (Taunton, Hassig, and Schreiber 1996). We next asked whether the processing of MEP-1 by DPF-3, or the lack thereof, might alter the composition of MEP-1-containing protein complexes, in addition to affecting its stability in *mep-1(T3E)* mutant animals. To address this, we immunoprecipitated MEP-1 from wild-type and *dpf-3Δ* mutant animals and analyzed the associated proteins by mass spectrometry. As a control for comparison, we also performed IP-MS on LIN-42 using the strain *lin-42(xe235[lin-42::3xFLAG::GFP::HiBit])* (kindly provided by Kathrin Braun).

As shown in Fig. 4B, Immunoprecipitation followed by mass spectrometry (IP-MS) of MEP-1 from wild-type animals revealed enrichment of its known interaction partner LET-418, as well as SPC-1 and F54A3.5, relative to LIN-42. However, the corresponding comparison between MEP-1 from *dpf-3Δ* mutant animals and LIN-42 did not reveal any additional interacting proteins or loss of canonical interactors (Suppl. Fig. 2B, please note that different FDR and S0 values are used for better visualization, see methods for details).

Moreover, a direct comparison between wild-type and *dpf-3Δ* MEP-1 IP-MS datasets did not identify any differentially enriched proteins (Fig. 4C), suggesting that the absence of processing by DPF-3 does not alter the composition of MEP-1 associated protein complexes.

### *mep-1(T3E)* mutant animals partially recapitulate the *dpf-3* loss of function phenotype

Studies have shown that *mep-1* plays an important role in *C. elegans* development, with its loss of function resulting in defective gonadogenesis and impaired oocyte production (Belfiore et al. 2002). The *mep-1* gene is essential for viability. The *mep-1(q660)* deletion mutant, which removes 635 amino acids, is not viable in the homozygous state but can be maintained in a heterozygous form (Belfiore et al. 2002). Also, RNAi mediated knockdown of *mep-1* results in lethality (Belfiore et al. 2002; Unhavaithaya et al. 2002).

Since we observed a significant decrease in the steady state levels of MEP-1 protein upon the *T3E* mutation, we sought to assess its effect on the physiology and the reproductive fitness of the animals. Using CRISPR/Cas9-mediated genome editing, we generated *mep-1(xe311[T3E])* (referred to as *mep-1(T3E)* hereafter; please note the *xe311* allele does not have an epitope). The *mep-1(T3E)* mutant animals were backcrossed three times to eliminate unintended off-target mutations. Unlike the *mep-1(q660)* deletion mutants, homozygous *mep-1(T3E)* animals could be maintained. These mutant animals appeared superficially wild-type but exhibited reproductive defects upon closer examination, such as unfertilized oocytes and defective embryos. Interestingly, loss of *dpf-3* function also results in reproductive defects, including reduced brood size (Gudipati et al. 2021). To quantify the reproductive defects in *mep-1(T3E)* mutant animals, we counted the number of offspring produced by culturing the animals at 25℃. As shown in Fig. 5A, *mep-1(T3E)* mutant animals displayed a significantly reduced brood size compared to wild-type animals mirroring the *dpf-3* loss-of-function mutant animals.

**Fig. 5.**
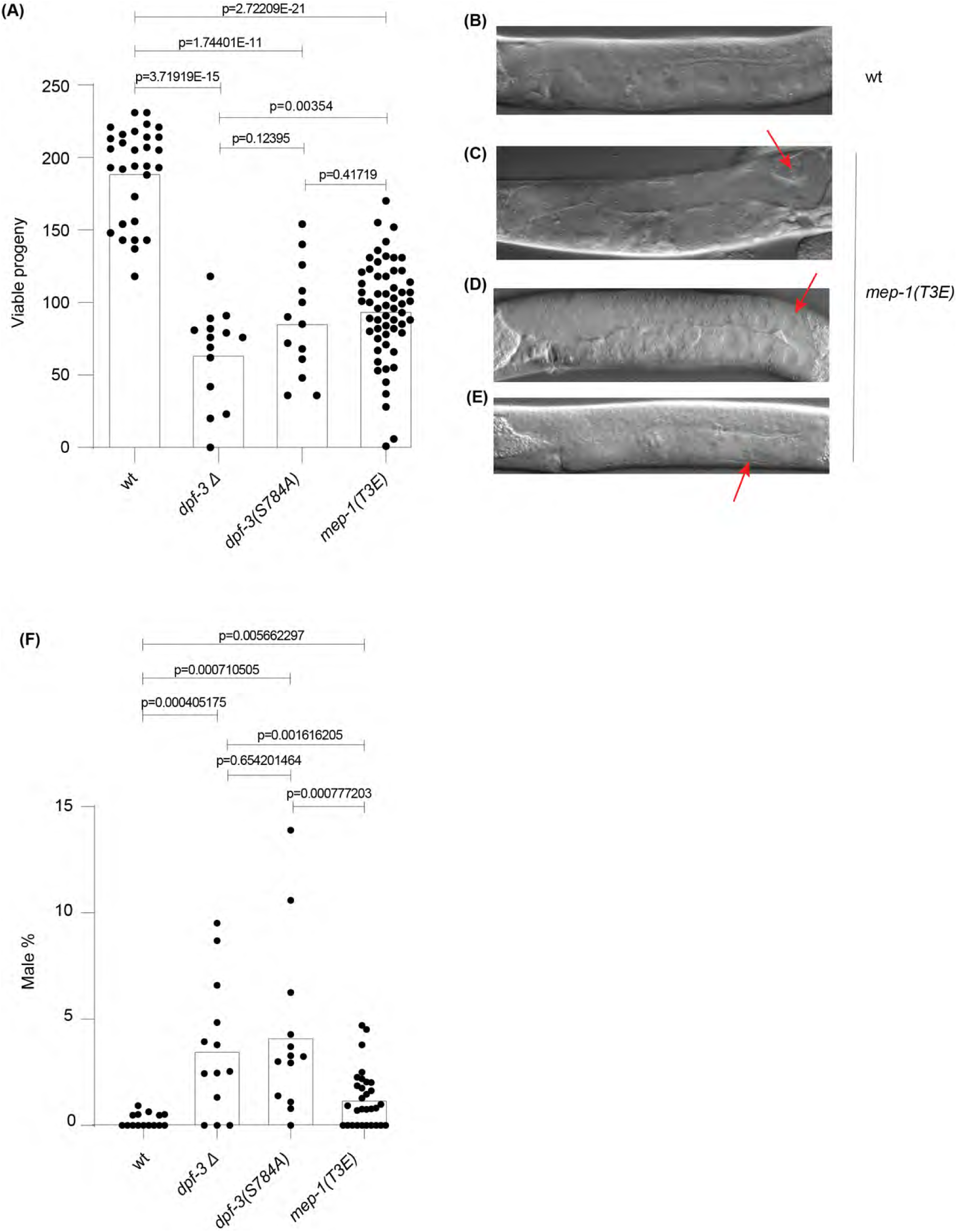
*mep-1(T3E)* mutant animals display reproductive defects (A) Box plot of the brood size (y-axis) of animals with different genetic background (x-axis) grown at 25℃. Each dot represents the total viable progeny of an individual animal. The height of the bar represents mean value. The decreased brood size in *dpf-3* loss-of-function mutant animals was already shown in (Gudipati et al. 2021), but was quantified again during the current work and shown for ease of comparison. (B-E). Representative Differential Interference Contrast (DIC) images of wild-type (B) and *mep-1(T3E)* (C-E) mutant animals grown at 25℃. Defective gonad migration (C) and premature oocyte development immediately after the bend (D), and oocytes that fail to develop properly and acquire a membrane (E) are indicated with red arrow. (F) Plot showing the percentage (%) of males (y-axis) in different genetic background (x-axis). Each dot represents the % of males in the total offspring of that particular worm. The height of the bar represents mean value.

To better understand the cause of reduced brood size in *mep-1(T3E)* mutant animals, we conducted a detailed microscopic analysis of their gonadal development and germline morphology in animals cultured at 25℃. Phenotypic defects in the germline were observed in over 70% of *mep-1(T3E)* mutant animals (n = 50), compared to approximately 15% of wild-type controls (n = 50). In *mep-1(T3E)* mutant animals, the most common abnormalities included premature oocyte formation (∼20%), enlarged oocytes (∼30%), and gonadal arm migration defects (∼25%) (The percentages refer to all the animals). Notably, many animals exhibited multiple concurrent defects, complicating classification in some cases. In wild-type animals, ∼10% displayed gonadal arm migration defects, and ∼5% showed enlarged oocytes (Fig. 5B-E), which is consistent with previous findings that elevated temperature impairs oogenesis in natural *C. elegans* variants (Petrella 2014).

Additionally, we observed a high incidence of males in *dpf-3* loss-of-function mutant animals compared to wild-type animals. While wild-type animals exhibited an average male frequency of 0.23%, *dpf-3Δ*, and *dpf-3(S784A)*, mutant animals displayed a significantly higher male frequency of 3.19% and 3.73%, respectively. Notably, *mep-1(T3E)* mutant animals also showed an elevated male frequency of 1.28%, partially mirroring the phenotype observed in *dpf-3* loss-of-function mutant animals (Fig. 5F).

## Discussion

Proteases play critical roles in maintaining cellular homeostasis and regulating physiological processes, and their dysfunction is often linked to disease states. The present study broadens our understanding of the substrate specificity and physiological relevance of DPF-3, a member of the DPPIV family proteases, by combining unbiased proteomic approaches and functional validation. Our findings reveal a previously unappreciated complexity in the substrate recognition and cleavage patterns of DPF-3 providing new insights into its role in nematode development and reproduction.

### Broad substrate specificity of DPF-3

Our study deepens the understanding of the DPPIV family proteases, particularly focusing on DPF-3 in *Caenorhabditis elegans.* The observation of substrate-product pairs in our TAILS dataset enabled the identification of novel substrates of DPF-3 *in vivo*. The differential abundance of substrates and products in wild-type and *dpf-3* mutant strains reflects both the efficiency of cleavage and the stability (or the detectability in mass spectrometer) of the resulting peptides. We have recently shown that DPF-3 tolerates amino acids other than Pro or Ala at the P1 position, including Thr *in vitro* (Gudipati et al. 2024). The identification and subsequent validation of MEP-1 as a substrate containing Thr at the P1 position provide the first *in vivo* physiological substrate of a DPPIV family member. These findings suggest that DPF-3-mediated processing can occur even when non-canonical amino acids are present at the P1 position, potentially expanding the biological relevance of this protease. The ability of DPF-3 to process substrates with Thr at the P1 position raises the possibility that human DPPIV family members may also have broader substrate repertoires than previously appreciated.

Despite fractionating the samples and attempting to achieve deep coverage of the proteome, we detected and quantified only a limited number of peptides representing ∼5000 proteins, about 1/4^th^ of the *C. elegans* proteome. We believe that increasing the proteome coverage by using different enzymes, such as Chymotrypsin, GluC, elastase or AspN (Dau, Bartolomucci, and Rappsilber 2020) for proteome digestion could help identify additional substrates. Comprehensive substrate profiling of these proteases, potentially leveraging unbiased approaches like TAILS, could provide critical insights into their physiological and pathological roles.

### DPF-3-mediated proteolysis of MEP-1: A crucial mechanism in reproductive development

Rendering MEP-1 resistant to DPF-3 cleavage disrupted oocyte development, impaired gonadal arm migration, and caused reproductive defects in *C. elegans*. These findings underscore the critical role of DPF-3 mediated proteolytic regulation of MEP-1 in ensuring proper reproductive development. This represents a novel layer of post-translational control that may be conserved across species, given the functional similarities between DPF-3 and its mammalian orthologs, DPP8/9.

DPF-3 appears to influence reproduction by targeting distinct substrates essential for male and female germline development. Our previous studies revealed that DPF-3 regulates the male germline by processing specific Argonaute proteins, WAGO-1 and WAGO-3. DPF-3-mediated maturation of these nascent proteins enhances their functional roles. Specifically, processing of WAGO-3 stabilizes its steady-state levels, while processing of WAGO-1 ensures the correct complement of 22G RNAs (siRNAs) is loaded. Loss of *dpf-3* function leads to de-silencing of transposable elements, DNA damage, and fully penetrant sterility due to defective spermatogenesis (Gudipati et al. 2021).

DPF-3 may regulate female germline function by targeting the essential protein MEP-1. Previous studies have shown that deletion or RNAi-mediated depletion of *mep-1* results in defective gonadal arm migration and impaired oogenesis (Belfiore et al., 2002). Notably, these defects were recapitulated in DPF-3 refractive *mep-1(T3E)* mutant animals, indicating the necessity of MEP-1 processing for its biological activity. Interestingly, *mep-1(T3E)* mutant animals exhibited a higher incidence of males, akin to *dpf-3* mutant animals, suggesting a role for DPF-3 in chromosome segregation and/or spermatogenesis. This phenotype may arise from disrupted processing of MEP-1 or other substrates critical for chromosome segregation or germline differentiation.

Despite the significant effect of DPF-3-mediated cleavage on MEP-1 stability, our IP-MS analyses indicate that the processing does not substantially alter the composition of MEP-1-associated protein complexes. This suggests that the proteolytic event primarily regulates substrate abundance rather than complex assembly or interaction specificity. Given that MEP-1 is part of chromatin remodelling complexes with LET-418 and HDA-1, DPF-3’s role might be to fine-tune the dosage of such regulatory proteins rather than remodel their interaction networks.

It is interesting to note that MEP-1 protein is more severely destabilized in DPF-3 refractive mutant background (*mep-1(T3E)*) than in the *dpf-3Δ* mutant background. The *C. elegans* DPPIV family is comprised of six potentially active members, *dpf-1* through *dpf-6*, and we speculate that in the absence of DPF-3 activity, other members of this family may partially compensate for the loss of function by cleaving shared substrates, including MEP-1. In contrast, in the *mep-1(T3E)* background, the cleavage site of MEP-1 is mutated, rendering it refractory to processing by DPF-3 and potentially all other DPPIV family proteases. This likely explains the pronounced reduction in MEP-1 protein abundance in the *mep-1(T3E)* mutant animals, underscoring the physiological importance of this non-canonical cleavage event.

Collectively, our results establish MEP-1 as a bona fide physiological substrate of DPF-3, and highlight that its processing is crucial for executing its biological functions. The implications of these findings are multifaceted. Firstly, our findings emphasize the importance of substrate profiling in elucidating the full scope of protease activity and function. The lack of comprehensive substrate data has been a significant bottleneck in understanding the mechanistic roles of DPPIV family proteases (Mulvihill and Drucker 2014). Innovative proteomic approaches, such as TAILS, could be extensively used and extended to other DPPIV family members to uncover novel substrates and processing events that could reveal unexpected biological functions. Secondly, this study establishes DPF-3 as a versatile protease with broader substrate specificity than previously recognized and underscores its critical role in reproductive development through the regulation of MEP-1. In conclusion, our study extends our understanding of DPPIV family proteases by identifying substrates with non-canonical P1 residues. These findings suggest the need for a more nuanced perspective on the catalytic capabilities of these enzymes, which could have broader implications.

### Limitations and future directions

Our studies rely on mass spectrometry for substrate identification, which might miss low abundance substrates. Additionally, the usage of trypsin for lysate digestion could be replaced by other enzymes that could generate a wide variety of peptides increasing the likelihood of peptide detection. Performing TAILS by using a different enzyme for proteome digestion could help in identifying additional substrates of DPF-3.

## Materials and Methods

### A. *elegans* strains and DNA oligonucleotides

DNA oligonucleotides used in this study are listed in Suppl. Table 3. The *C. elegans* strains used in this study are listed in Suppl. Table 4.

### Maintenance of *C. elegans* and worm extract preparation

The animals were grown on Nematode Growth Medium (NGM) 2% with *Escherichia coli* NA22 bacteria as a food source at 25℃. To synchronize the worms, we extracted the eggs from gravid adults by bleaching solution (30% (v/v) sodium hypochlorite, 5% chlorine (Thermo Fisher Scientific; 419550010), 0.75M KOH)). The eggs were left to hatch in M9 buffer (42 mM Na_2_HPO_4_, 22 mM KH_2_PO_4_, 86 mM NaCl, 1 mM MgSO_4_) without food for 12-16 hrs, generating synchronized, arrested L1 stage worms. The staged L1 worms were plated on NGM 2% plate containing NA22 bacteria as a food source and were collected at the indicated developmental stages. The worms were washed three times in M9 buffer and transferred to 2 ml tubes (Sarstedt, Ref: 72.693.005) with Zirconia/Silica beads (Biospec, Cat. No. 110791052) in lysis buffer (50mM Tris-HCl, pH 7.5, 150 mM NaCl, 1% TRITON X-100, 1 mM EDTA) supplemented with 1mM PMSF and 1 tablet of cOmplete Protease inhibitor (Roche, cat. No. 11 873 580 001) per 50 ml. The worms were lysed using the MP Biomedicals Fast Prep-24 5G bead beater for 16 cycles at 8m/Sec. The lysate was centrifuged at 16,100 g for 10 min at 4℃ and the supernatant was transferred to a fresh tube and the protein concentration was measured using standard bradford assay.

### Construction of transgenic animals

Transgenic lines were generated using CRISPR-Cas9 genome editing technology as described in (Katic, Xu, and Ciosk 2015; Arribere et al. 2014). Not1 linearized pIK198 was used to clone the sgRNAs by Gibson assembly. A mix containing 50 ng/µl pIK155, 10 ng/μl of pIK208, 100 ng/μl of pIK198 with the cloned sgRNA, 20 ng/µl of AF-ZF-827 primer, 20 ng/µl of single stranded primer as a template for homologous recombination targeting the gene of interest was injected into Bristol N2 worms unless indicated otherwise. Transgenic worms were isolated using *dpy-10* co-conversion method as described in (Arribere et al. 2014) For an alternative RNP-based editing strategy, ribonucleoprotein complexes (RNPs) were prepared by first combining the following components in the stated order: Cas9 protein (∼30 pmol), tracrRNA (∼90 pmol), crRNA 4 µg/µL; ∼95 pmol total. The mixture was gently pipetted and incubated at 37 °C for 15 minutes to allow efficient RNP complex formation.

Following RNP assembly, the following components were added to the reaction: ssODN donor (1 µg/µL), pRF4(*rol-6*)(su1006) plasmid (500 ng/µL) and pIK127 (eft-3::GFP) to a final concentration 20ng/ul used as a co-injection marker. The final mix was gently pipetted to ensure homogeneity before injection into. The transgenic lines were confirmed by PCR and outcrossed to wild-type N2 at least three times. All the sgRNA, crRNA sequences and primers used for introducing desired changes are listed in Suppl. Table 3. The endogenously tagged *mep-1* strain, (*mep-1(xe312[3xflag::linker::mep-1])*IV) has a 3x *flag-linker* sequence between the 52^nd^ and 53^rd^ amino acids. Endogenously tagged *mep-1(T3E)* strain, (*mep-1(xe311aga19[3xflag::linker::mep-1(T3E)*])IV was obtained by inserting 3x *flag-linker* sequence between the 52^nd^ and 53^rd^ amino acids into *xe311* strain.

### Terminal Amine Isotopic Labeling of Substrates (TAILS)

The worms were collected in ice cold PBS and washed three times with PBS. Another wash was done with PBS containing 5mM DTT and 1mM PMSF. A final wash was done with lysis buffer (PBS with Guanidine HCl 6M; DTT 5mM and 1 tablet of cOmplete Protease inhibitor (Roche, cat. No. 11 873 580 001) per 50 ml)) followed by resuspension in 500 µl of lysis buffer and snap freezing in liquid nitrogen. The worms were disrupted (5 × 25 seconds) using 0.5 mm Zirconia/Silica beads (Biospec, Cat. No. 110791052) at intensity 8m/s^2^ in an MP Biomedicals Fast Prep-24 5G bead beater. The pH was adjusted to 8.5 using 1% ammonium hydroxide. The lysate was boiled at 65℃ for 30 min and spun down for 10 min at 14000 rpm at 4℃. The supernatant was transferred to a fresh tube and protein concentration was measured by Bradford assay. 250 µg of protein per sample was used in the TAILS protocol that was adapted from (Kleifeld et al. 2011) and is described below.

### Protein precipitation to remove buffer salts, amine scavengers

Four times the sample volume of freezer-cold methanol was added to the sample, followed by mixing and the addition of two times the sample volume of freezer-cold chloroform with further mixing. Three times the sample volume of water was added and mixed followed by centrifugation for 2 min at 4℃ and 5000 rcf. The protein precipitate was clearly visible at the interface of chloroform (bottom) and aqueous (top) phases. The liquid was carefully removed and 750 µl of methanol was added, mixed, and the pellet was crushed with a pipet tip. The sample centrifuged at 6,500 rcf for 15 minutes. The protein pellet was washed twice with methanol.

### Protein re-solubilization, reduction/alkylation and TMT labeling

The protein pellet was air-dried, then 30 µl of 6M GuHCl in 200 mM HEPES pH 8 was added, followed by 70 µl of 200 mM HEPES pH 8 and 11 µl of 100 mM TCEP in 200 mM HEPES to obtain a final concentration of 10 mM TCEP and 200 mM HEPES. After 1 hr of incubation while shaking, 13 µl of 250 mM iodoacetamide was added and incubated while shaking for 30 min in the dark. Later, 800 µg of TMT 6plex reagent stock (Thermo Fisher Scientific 126-131, Lot# PG202321C) in 124 µl anhydrous DMSO was added to every sample (to give 50% DMSO final concentration) and incubated while shaking for 1 hr. Then 25 µl of 1 M ethanolamine (30 µl 99.9% ethanolamine + 470 µl 200 mM HEPES) was added to give 0.5 % final concentration and incubated while shaking for 30 min to quench unreacted TMT labels. Six quenched protein samples were combined to a final volume of 1.638 ml, into two different 15 ml Falcon tubes.

### Protein precipitation and digestion

To each Falcon tube, 8x the sample volume of freezer-cold acetone and 1x the sample volume (1.638 mL) of freezer-cold methanol were added, vortexed, and protein was allowed to precipitate at −20℃ overnight. After centrifugation at 4℃ for 15 min at 7,100 rcf, the supernatant was carefully removed and discarded. 5 ml of freezer-cold methanol was used to wash the pellet. 23 µl of 100 mM NaOH was added, then 75 µl of 1M HEPES pH 8, followed by 1.402 ml of water to obtain a final concentration of 1 mg/ml of protein in a total volume of 1.5 ml 50 mM HEPES, which was transferred to a 2-ml Eppendorf tube. To dissolve residual pelleted material, Falcon tubes were washed once with 100 µl of 50 mM HEPES and the material transferred to the same Eppendorf tube. Fifteen µg (75 µl, 0.2 µg/µl) of Promega sequencing-grade trypsin in Promega trypsin resuspension solution was added (protease to protein ratio of 1:100), mixed and incubated overnight at 37℃.

### Desalting and preparation of proteome (pre-TAILS) samples for LC-MS single-shot analysis and, peptide fractionation

100 µl (100 µg) of TMT 6plex-labeled protein digests were desalted on a 50 mg SepPak cartridge, eluted with 500 µl of 50% acetonitrile, 0.1% TFA. Desalted peptides were dried in a speedvac (ca. 10 µg (50 µl) in a glass insert for single-shot quality control LC-MS analysis, and the remaining (approximately 90 ug) in an Eppendorf tube for subsequent peptide fractionation.

### Negative selection using HPG-ALDII polymer

Two 10-kDa MWCO Amicon columns (0.5 m; centrifugal filter Amicon ultra 0.5 ml) were washed with 450 µl of 100 mM NaOH at 14k rcf for 10 min, then 3x washed with 450 µl of water, then loaded with 100 µl (maximum recommended amount) of HPG-ALDII polymer (60 mg/ml, purchased from The University of British Columbia, Canada), topped off with water and centrifuged at 14k rcf for 5 min, then washed with 450 µl of water at 14k rcf for 10 minutes. The filter was inverted and the washed polymer slurry was centrifuged into the remaining 1.4 ml of TMT 6plex-labeled protein digest (polymer to peptide ratio of 5:1, w/w), immediately followed by the addition of 6 µl of 5M NaCNBH_3_ (20 mM final concentration) and 10 µl of 2M HCl to obtain a pH of 6, then incubated with shaking at 37℃ overnight.

### Recovery of the TMT 6plex-labeled N-terminome (post-TAILS sample)

160 µl of 1 M Tris HCL pH 6.8 was added to the peptide-polymer mixture (final concentration of 100 mM Tris, pH 6.5), incubated in shaker at 37℃ for 30 minutes. Two 10-kDa MWCO Amicon columns were pre-washed with 400 µl of 100 mM NaOH, centrifuged at 10k rcf for 10 min, the flow-through was discarded. Then 400 µl of water was loaded, centrifuged at 10k rcf for 10 min, the flow-through was discarded. This step was repeated 3 times. The mixture was loaded, centrifuged at 10k rcf for 5 min and collected into a 2 ml tube. The flow-through was transferred to a fresh 1.5 ml Eppendorf tube. The polymer was washed with 100 µl of water and all liquid (including polymer) was transferred to a fresh tube. The filter was inverted, and whatever liquid might have been left was spun into the 1.5 ml tube with all the combined flow-through.

### Desalting and preparation of post-TAILS sample for LC-MS single-shot analysis and peptide fractionation

The pooled sample was acidified with 50 µl of 5% TFA (0.2% final concentration) to obtain a pH of 2-3, and then stored at −20℃. The samples were desalted on 50 mg SepPak tC18 cartridges, eluted with 500 µl of 50% acetonitrile 0.15% TFA. 10% of the eluate was transferred into an autosampler glass vial, dried in a speedvac and prepared for single-shot quality control LC-MS analyses on an Orbitrap Fusion Lumos MS, and 90% of the eluate was dried and prepared for further chromatographic fractionation.

### High pH reversed phase chromatographic peptide fractionation

To fractionate TMT-labeled peptides, aliquots of 55 μg of the dried pre-TAILS sample, or 100 μg of the post-TAILS sample were dissolved in 20 μL of 10 mM ammonium formate (pH 10, Buffer A). High-pH reversed-phase chromatography was then employed using an Agilent 1100 HPLC system with an autosampler, a YMC Triart C18 0.5 x 250 mm column, and a fraction collector. The peptides were separated using the following binary buffer system: high-pH Buffer A (20 mM ammonium formate in water, pH 10) and high-pH Buffer B (20 mM ammonium formate pH 10 in 90% acetonitrile) at a flow rate of 12 μL/minute. The gradient duration was a total of 115 min, with mobile phase compositions in %B as follows: 0-5 min at 2%-15%, 5-15 min at 15%-25%, 15-80 min at 25%-45%, 80-85 min at 45%-65%, 85-95 min at 65%-100%, 95-100 min at 100%, 100-101 min at 100%-2%, and 101-115 min at 2%. After collecting 72 fractions from 3-112 min, they were combined (concatenated) to 24 super-fractions (Wang et al. 2011).

### LC-MS analysis of TMT labeled peptide samples

Proteome (pre-TAILS) fractions were analyzed in 110 min and N-terminome (post-TAILS) fractions were analyzed in 85 min runs using an Orbitrap Fusion Lumos mass spectrometer. To prepare for each LC-MS run, about 1 μg of peptides were loaded onto a PepMap 100 C18 2 cm trap using an EASY nLC-1000 system. On-line peptide separation was achieved using a 15 cm EASY-Spray C18 column with a linear gradient of increasing acetonitrile concentration in 0.1% formic acid and water, flowing at a rate of 250 nL/minute. An Orbitrap Fusion Lumos Tribrid mass spectrometer was operated in a data-dependent mode to quantify TMT reporter ions using synchronous precursor selection-based MS3 fragmentation, as described in (McAlister et al. 2014). Briefly, every 8 seconds, the most intense precursor ions from the Orbitrap survey scan (MS1) were selected for collision-induced dissociation fragmentation with a normalized collision energy set to 35%. The MS2 CID spectrum was generated by the ion-trap analyzer from which the 10 most abundant notches for the MS3 scan were selected. The MS3 spectrum was then recorded using the Orbitrap analyzer at a resolution of 15,000.

### Data analysis

Quality control of TMT protein labeling efficiency, digestion, depletion, potential N-terminal enrichment and TMT-reporter ion-based peptide and protein quantification etc., were performed with Proteome Discoverer PD 2.5 (Thermo Scientific) and the einprot R package (Soneson et al. 2023). MS2 fragment ion spectra were searched in PD with the Sequest HT search engine in Proteome Discoverer against the *Caenorhabditis elegans* Uniprot protein fasta database (query: organism:“Caenorhabditis elegans [6239]” AND proteome:up000001940, canonical sequences & isoforms), downloaded on May 03 2021, as well as a database that contains commonly observed contaminants (a combination of the contaminant database included with the MaxQuant software, the common Repository of Adventitious Proteins, cRAP, and FMI-specific tags and QC sample sequences). A maximum of one missed cleavage was tolerated for Arg-C (semi) enzymatic digestion specificity. Fixed peptide modifications were set for Carbamidomethyl / +57.021 Da (C), TMT 6plex / +229.163 Da (K); and variable peptide modifications were allowed for Oxidation / +15.995 Da (M), Deamidated / +0.984 Da (N), Acetyl / +42.011 Da (N-Terminus), TMT6plex / +229.163 Da (N-Terminus). Peptide-to-spectrum matches (PSMs) were validated using the target-decoy search strategy (Elias and Gygi 2007) and Percolator with a strict confidence threshold of 0.01, and a relaxed confidence threshold of 0.05. Unique+razor peptides were used for MS3-based TMT 6plex reporter-ion quantification, taking into account quan value correction for isotopic impurities of the TMT reagents (Lot# PG202321C), with the requirements of maximum MS1 co-isolation threshold 50%, and minimum PSP mass matches of 65%. Median centering normalization and MinProb imputation were applied using the PTM workflow in the in-house developed einprot R package, version 0.6.8. For the protein analysis, only Master Proteins were considered. Figures were generated using einprot results and in-house R scripts. TAILS data can be downloaded from PRIDE with accession number PXD042290.

### *In vitro* protease assay by mass spectrometry

All the synthetic peptides were purchased from Thermo Fisher Scientific, and their cleavage by recombinant enzymes was analyzed as described in (Gudipati et al. 2021). Briefly, the lyophilized peptides were dissolved in TBS buffer supplemented with 2 mM TCEP to obtain a stock solution with a final concentration of 0.1 nmol/µl. One µg of either wild-type or *S784A* mutated DPF-3 protein and each peptide at a final concentration of 1 pmol/µl were used in reactions at a final volume of 100 µl. The assay was performed at 21°C and aliquots (20 µl) were taken at the indicated times and mixed with 180 µl of 0.1 % trifluoroacetic acid, 2 mM TCEP, 2 % acetonitrile in water to stop the reaction.

The peptides were analyzed by capillary liquid chromatography tandem mass spectrometry (LC-MS). Peptides were loaded onto a 75 μm×15 cm ES811 column (Accucore C4, 2.6 μm, 150 Å) at a constant pressure of 800 bar, using an EASY-nLC 1000 liquid chromatograph with one-column set up (Thermo Scientific). Peptides were separated at a flow rate of 300 nl/min with a linear gradient of 2-6 % buffer B in buffer A in 2 min, followed by a linear increase from 6 %-22 % in 20 min, 22 %-28% in 4 min, 28 %-36 % in 2 min, 36% −80 % in 1 min and the column was finally washed for 5 min at 90% buffer B (buffer A: 0.1% formic acid in water; buffer B: 0.1% formic acid in acetonitrile). The column was mounted on a DPV ion source (New Objective) connected to an Orbitrap Fusion mass spectrometer (Thermo Scientific), data were acquired using 120,000 resolution, for the peptide measurements in the Orbitrap and a top T (1.5 s) method with HCD fragmentation for each precursor and fragment measurement in the ion trap following the manufacturer guidelines (Thermo Scientific). MS1 signals were quantified using Skyline 4.1 (MacLean et al. 2010).

### 3xFLAG-MEP-1 immunoprecipitation

The immunoprecipitation (IP) of proteins of interest was performed exactly as described in (Gudipati et al. 2021). Briefly, the anti-flag M2 Magnetic Beads (Sigma. Catalog Number M8823) were pre-washed thrice with TBS buffer (50 mM Tris-HCl pH 7.5, 150 mM NaCl). The cleared extract from total worms was added to 50 µl of 50 % bead suspension and incubated on a rotating wheel at 4℃ for 2 Hours. The beads were then washed once with lysis buffer that contained protease inhibitors and then washed 6 more times with TBS buffer without protease inhibitors and detergents.

### Mapping the *in vivo* cleavage sites and characterization of the protein complexes of MEP-1

Upon IP as described above, 5 µl of digestion buffer (3 M GuaHCl (Guanidine Hydrochloride), 20 mM EPPS (4-(2-Hydroxyethyl)-1-piperazinepropanesulfonic acid) pH 8.5, 10 mM CAA (2-Chloroacetamide), 5 mM TCEP (Tris(2-carboxyethyl)phosphine hydrochloride)) and 1 µl of Lys-C (0.2 µg/µl in 50 mM Hepes, pH 8.5) was added to the beads for incubation at 21°C for 2-4 h while shaking. Subsequently, 17 µl of 50 mM HEPES, pH 8.5 and 1 µl of 0.2 µg/µl trypsin was added and incubation continued at 37°C overnight. In the morning, another 1 µl of trypsin was added and incubation continued for 4 more hours.

Peptides of immunoprecipitated MEP-1 were analyzed by LC-MS/MS, essentially as described (Ostapcuk et al. 2018). In short, the peptides were separated with an EASY-nLC 1000 on a 50 μm × 15 cm ES801 C18, 2 μm, 100 Å column (Thermo Scientific) mounted on a DPV ion source (New Objective). They were measured with an Orbitrap Fusion (Thermo Scientific) using a top T (3 s) method as recommended by the manufacturer (Thermo Scientific). Andromeda implemented in MaxQuant (version: 1.5.3.8) (Cox et al. 2011) was used to search the CAEEL subset of the UniProt (version: 2017_04) combined with the contaminant database from MaxQuant and label-free quantification (LFQ) (Cox et al. 2014) was used with a protein and peptide FDR of 0.01. Statistical analysis was done in Perseus (version: 1.5.2.6) (Hubner et al. 2010; Tyanova et al. 2016). MS1 signals of selected MEP-1 peptides were quantified using Skyline 4.1 (MacLean et al. 2010) (Fig. 3D).

### Characterizing the protein complexes of MEP-1

Immunoprecipitated MEP-1(bait) from *flag::mep-1* and *dpf-3Δ;mep-1::flag* and LIN-42 (control) were analyzed by LC-MS/MS as described above. Raw mass spectrometry (MS) data was processed using MaxQuant software (version 2.6.8.0) with the Andromeda search engine. Protein identification was performed against the *Caenorhabditis elegans* UniProt database (release 2025_05). Default MaxQuant parameters were used unless otherwise stated. The following key settings were applied: enzyme specificity was set to trypsin/P with a maximum of two missed cleavages; fixed modification was carbamidomethylation of cysteine, and variable modifications included methionine oxidation and N-terminal acetylation. The false discovery rate (FDR) was set to 1% at both the peptide-spectrum match (PSM) and protein levels. The minimum peptide length was set to 7 amino acids. Label-Free Quantification (LFQ) was enabled, with the minimum ratio count set to 2. The resulting proteinGroups.txt output was further analyzed in Perseus (version 2.011.0). Protein groups flagged as “Reverse”, “Only identified by site”, or “Potential contaminant” were removed from the dataset. LFQ intensities were log2-transformed to approximate normal distribution. Missing values were imputed from a normal distribution (width = 0.3, downshift = 1.8) to simulate signals of low-abundance proteins. For comparative analysis, categorical annotation was added to distinguish among the experimental groups: MEP-1, *dpf-3Δ;*MEP-1, and LIN-42. Two-tailed Student’s t-tests were performed for the following pairwise comparisons: For the MEP-1 *vs*. LIN-42 and MEP-1 *vs*. *dpf-3Δ;*MEP-1 comparisons, significance thresholds were defined using a permutation-based FDR of 0.05 and S0 = 2, balancing stringency and detection sensitivity. However, for the *dpf-3Δ;*MEP-1 *vs*. LIN-42 comparison, initial statistical parameters failed to separate bait and control in the resulting volcano plot due to low signal dispersion and overlap in protein abundance. To resolve this, a more permissive threshold (FDR = 0.1, S0 = 1.22) was empirically selected. This adjustment allowed the visualization of significantly enriched proteins in both bait and control samples, providing a more informative interpretation of the differential interaction landscape.

### Brood size counting

L4 hermaphrodites, designated as P0s, were individually placed on 35-mm plates seeded with *E. coli* OP50 and incubated at the specified temperature. Once enough eggs had been laid, each P0 was transferred to a fresh plate, repeating this process until egg-laying ceased. The number of viable F1 progeny was recorded, and results were displayed as a boxplot, with each dot representing the total viable progeny from a single P0.

### RNA quantification by RT-qPCR

Synchronized animals grown at 25℃ on NGM plates were collected at the L4 stage in biological triplicates. The strains used are: *flag::mep-1*, *dpf-3Δ*; *flag::mep-1* and *flag::mep-1(t3e).* Following washes, 1 ml of TRIzol (Thermo Fisher Scientific) was added and frozen at −80°C. The animals were thawed and incubated at 65°C for 30 minutes, followed by addition of 200 ul of chloroform and centrifugation (12,000 × g, 15 min, 4℃). The upper aqueous layer was mixed with 100% ethanol and GlycoBlue™, incubated at 4℃ for 1 h, and centrifuged to pellet RNA. After washing with 70% ethanol and air-drying, RNA was resuspended in nuclease-free water. RNA concentration and purity were assessed using a NanoDrop spectrophotometer. For cDNA synthesis, 1 µg of total RNA was reverse transcribed using the Thermo Scientific RevertAid First Strand cDNA Synthesis Kit according to the manufacturer’s instructions. Quantitative PCR (qPCR) was performed using SYBR Green Master Mix (Applied Biosystems) on a real-time PCR system. Reactions were run in triplicate for each sample. Expression levels were normalized to a housekeeping gene, and relative quantification was calculated using the ΔΔCt method.

### Image acquisition

The animals were mounted on 2% (w/v) agarose pads with 10 μl of 10 mM levamisole. Images were captured on a Nikon eclipse Ni microscope equipped with DS-QI2 camera using NIS elements BR 5.42.04 software. Differential Interference Contrast (DIC) images were acquired with a 40x oil immersion objective. Fiji software (Schindelin et al. 2012) was used for image processing and selection of representative images.

## Acknowledgements

We thank Kathrin Braun for the gift of epitope-tagged *lin-42* transgenic animals. We thank Iskra Katic and Anca Neagu for technical support in the generation of other transgenic animals. We thank Dimosthenis Gaidatzis for his input in analyzing the TAILS data and Thomas Welte for his inputs and stimulating discussions. We are grateful to Vytautas Iesmantavicius for help with peptide fractionation and recording of LC-MS data. Some strains were provided by the *Caenorhabditis* Genetics Center (CGC), which is funded by NIH Office of Research Infrastructure Programs (P40 OD010440).

This work was supported by by the National Science Center (NCN), Poland, SONATA BIS 2021/42/E/NZ1/00336 and OPUS 2022/45/B/NZ2/02183 (to R.K.G) and the NCCR RNA & Disease, a National Centre of Excellence in Research, funded by the Swiss National Science Foundation (grant number 182880), by the Novartis Research Foundation through the Friedrich Miescher Institute for Biomedical Research (to H.G).

## Author Contributions

RKG and HG conceived the project, acquired the funds, designed and analyzed experiments and wrote the manuscript with inputs from the rest of the authors. IA generated one of the strains, performed western blot and RT-qPCR experiments, and analyzed mass spectrometry data to identify MEP-1 associated proteins. IA and AA characterized the *mep-1* mutant animals and quantified the brood size and *him* phenotype. JS digested the worm total lysate with trypsin, enriched and purified the peptides, and performed mass spectrometry for the TAILS experiments and analyzed the data using software developed by CS with input from Dimosthenis Gaidatzis. DH did the mass spectrometry for protease assays on specific synthetic peptides and MEP-1 immunoprecipitation.

## Data and material availability

The TAILS mass spectrometry proteomics data has been deposited at the ProteomeXchange Consortium via the PRIDE (Perez-Riverol et al. 2022) partner repository with the dataset identifier PXD042290 at https://www.ebi.ac.uk/pride/login. (Username, Password: 0pdipqjK). The MEP-1 IP-MS data is also deposited at ProteomeXchange Consortium via the PRIDE with the dataset identifier PXD065105 at http://www.ebi.ac.uk/pride (Username: Password: HKB6KqnWGHAJ). Published research reagents from the FMI are shared with the academic community under a Material Transfer Agreement (MTA) having terms and conditions corresponding to those of the UBMTA (Uniform Biological Material Transfer Agreement). The einprot package is available from GitHub (https://github.com/fmicompbio/einprot). Archives of the versions used here are available from https://doi.org/10.5281/zenodo.8031403 (v0.6.8) and https://doi.org/10.5281/zenodo.8031408 (v0.7.0).

## Supplementary Data

Suppl. Table 1: Peptides detected in TAILS experiments

Suppl. Table 2: Potential DPF-3 substrates as identified in TAILS.

Suppl. Table 3: Primers used in this study.

Suppl. Table 4: *C. elegans* strains in this study.

---

**Supp. Fig. 1.**
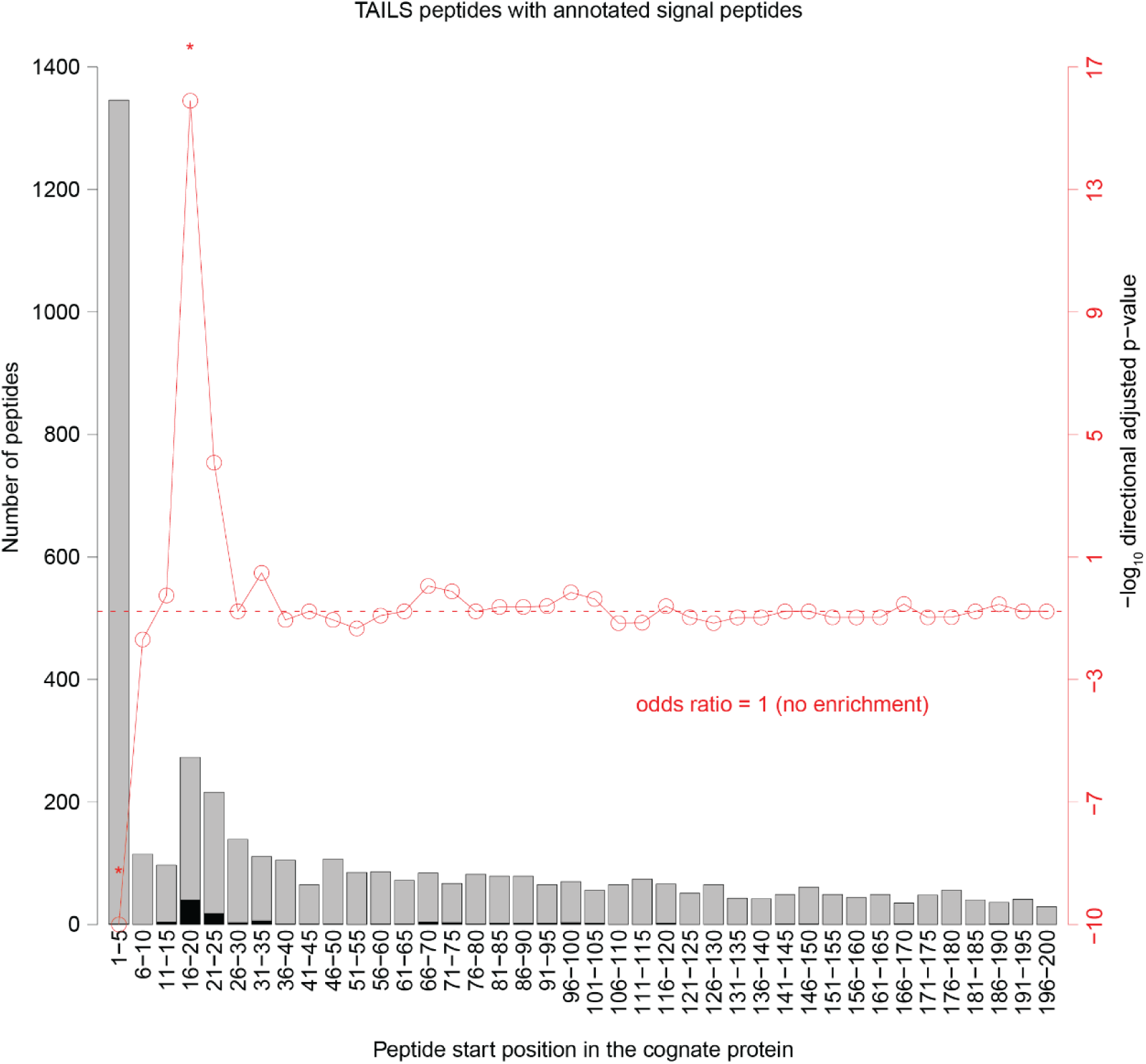
TAILS enriches protein N-termini Starting positions of peptides identified by TAILS are shown in 5 amino acid bins relative to the annotated start site for peptides originating within the first 200 amino acids. Peptides derived from proteins with an annotated signal peptide are indicated in black. The negative log_10_ directional, adjusted p-values (Fisher’s exact test) testing for the enrichment of signal peptides in proteins at a given bin position are shown in red. Null hypothesis: the fraction of signal peptide annotations is independent of the start position (odds ratio = 1). There is a significant under representation of annotated signal peptides in the TAILS peptide position bins 1-5, and an over representation between 16 and 25 (red asterisks)

**Suppl. Fig. 2.**
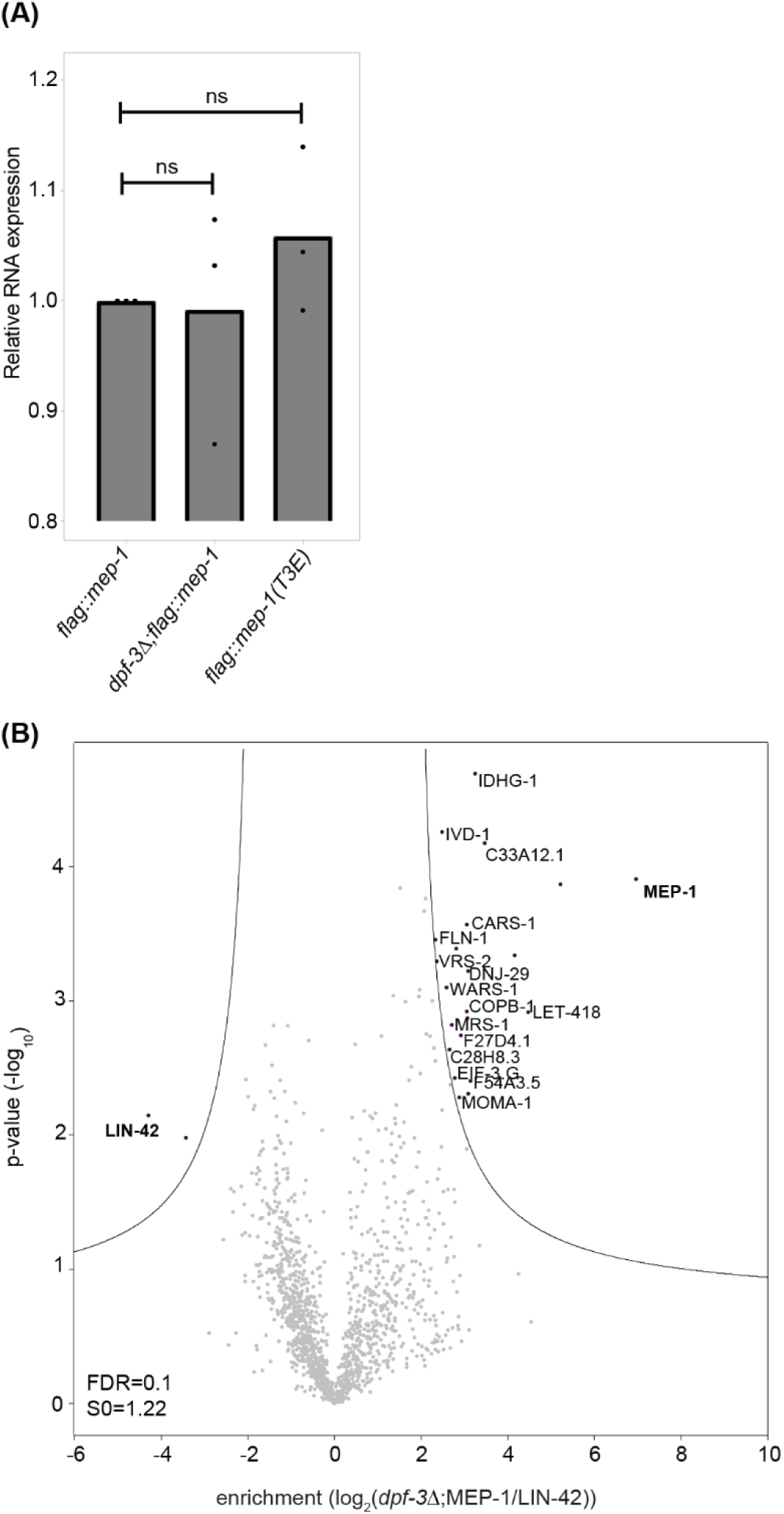
*mep-1* transcript levels are unaffected by *dpf-3*, and MEP-1 interactors are unchanged in *dpf-3Δ* mutant animals. A. RT-qPCR quantification of *mep-1* mRNA in wild-type, *dpf-3Δ*, and *mep-1(t3e)* mutant animals showing no significant differences in transcript abundance. B. IP-MS of MEP-1. The x-axis indicates the fold enrichment of the bait protein (MEP-1 from *dpf-3Δ* animals) and its interacting partners over control (LIN-42). Each dot represents a protein, the non-significant and the significant proteins are colored gray and black, respectively. False Discovery Rate (FDR) of 0.1 was used. The y-axis of the volcano plot indicates the p-values that were calculated by two-tailed Student t-test.

## References

Arribere, J. A., R. T. Bell, B. X. Fu, K. L. Artiles, P. S. Hartman, and A. Z. Fire. 2014. ‘Efficient marker-free recovery of custom genetic modifications with CRISPR/Cas9 in Caenorhabditis elegans’, Genetics, 198: 837–46.

Belfiore, M., L. D. Mathies, P. Pugnale, G. Moulder, R. Barstead, J. Kimble, and A. Puoti. 2002. ’The MEP-1 zinc-finger protein acts with MOG DEAH box proteins to control gene expression via the fem-3 3’ untranslated region in Caenorhabditis elegans’, RNA, 8: 725–39.

Cox, J., M. Y. Hein, C. A. Luber, I. Paron, N. Nagaraj, and M. Mann. 2014. ‘Accurate proteome-wide label-free quantification by delayed normalization and maximal peptide ratio extraction, termed MaxLFQ’, Mol Cell Proteomics, 13: 2513–26.

Cox, J., N. Neuhauser, A. Michalski, R. A. Scheltema, J. V. Olsen, and M. Mann. 2011. ‘Andromeda: a peptide search engine integrated into the MaxQuant environment’, J Proteome Res, 10: 1794–805.

Dau, T., G. Bartolomucci, and J. Rappsilber. 2020. ‘Proteomics Using Protease Alternatives to Trypsin Benefits from Sequential Digestion with Trypsin’, Anal Chem, 92: 9523–27.

Deacon, C. F. 2019. ‘Physiology and Pharmacology of DPP-4 in Glucose Homeostasis and the Treatment of Type 2 Diabetes’, Front Endocrinol (Lausanne), 10: 80.

Demuth, H. U., C. H. McIntosh, and R. A. Pederson. 2005. ‘Type 2 diabetes--therapy with dipeptidyl peptidase IV inhibitors’, Biochim Biophys Acta, 1751: 33–44.

Drag, M., and G. S. Salvesen. 2010. ‘Emerging principles in protease-based drug discovery’, Nat Rev Drug Discov, 9: 690–701.

Elias, J. E., and S. P. Gygi. 2007. ‘Target-decoy search strategy for increased confidence in large-scale protein identifications by mass spectrometry’, Nat Methods, 4: 207–14.

Gudipati, R. K., K. Braun, F. Gypas, D. Hess, J. Schreier, S. H. Carl, R. F. Ketting, and H. Grosshans. 2021. ‘Protease-mediated processing of Argonaute proteins controls small RNA association’, Mol Cell, 81: 2388–402 e8.

Gudipati, R. K., D. Gaidatzis, J. Seebacher, S. Muehlhaeusser, G. Kempf, S. Cavadini, D. Hess, C. Soneson, and H. Grosshans. 2024. ‘Deep quantification of substrate turnover defines protease subsite cooperativity’, Mol Syst Biol.

Hama, T., M. Okada, K. Kojima, T. Kato, M. Matsuyama, and T. Nagatsu. 1982. ‘Purification of dipeptidyl-aminopeptidase IV from human kidney by anti dipeptidyl-aminopeptidase IV affinity chromatography’, Mol Cell Biochem, 43: 35–42.

Harvey, L. M., P. M. Frederick, R. K. Gudipati, P. Michaud, F. Houle, D. Young, C. Desbiens, S. Ladouceur, A. Dufour, H. Grosshans, and M. J. Simard. 2025. ‘Dipeptidyl peptidase DPF-3 is a gatekeeper of microRNA Argonaute compensation in animals’, Nat Commun, 16: 2738.

Heymann, E., and R. Mentlein. 1978. ‘Liver dipeptidyl aminopeptidase IV hydrolyzes substance P’, FEBS Lett, 91: 360–4.

Hopsu-Havu, V. K., and G. G. Glenner. 1966. ‘A new dipeptide naphthylamidase hydrolyzing glycyl-prolyl-beta-naphthylamide’, Histochemie, 7: 197–201.

Hubner, N. C., A. W. Bird, J. Cox, B. Splettstoesser, P. Bandilla, I. Poser, A. Hyman, and M. Mann. 2010. ‘Quantitative proteomics combined with BAC TransgeneOmics reveals in vivo protein interactions’, J Cell Biol, 189: 739–54.

Johnson, D. C., C. Y. Taabazuing, M. C. Okondo, A. J. Chui, S. D. Rao, F. C. Brown, C. Reed, E. Peguero, E. de Stanchina, A. Kentsis, and D. A. Bachovchin. 2018. ‘DPP8/DPP9 inhibitor-induced pyroptosis for treatment of acute myeloid leukemia’, Nat Med, 24: 1151–56.

Katic, I., L. Xu, and R. Ciosk. 2015. ‘CRISPR/Cas9 Genome Editing in Caenorhabditis elegans: Evaluation of Templates for Homology-Mediated Repair and Knock-Ins by Homology-Independent DNA Repair’, G3 (Bethesda), 5: 1649–56.

Keane, F. M., N. A. Nadvi, T. W. Yao, and M. D. Gorrell. 2011. ‘Neuropeptide Y, B-type natriuretic peptide, substance P and peptide YY are novel substrates of fibroblast activation protein-alpha’, FEBS J, 278: 1316–32.

Kleifeld, O., A. Doucet, A. Prudova, U. auf dem Keller, M. Gioia, J. N. Kizhakkedathu, and C. M. Overall. 2011. ’Identifying and quantifying proteolytic events and the natural N terminome by terminal amine isotopic labeling of substrates’, Nat Protoc, 6: 1578–611.

Komatsu, R., T. Matsuyama, M. Namba, N. Watanabe, H. Itoh, N. Kono, and S. Tarui. 1989. ‘Glucagonostatic and insulinotropic action of glucagonlike peptide I-(7-36)-amide’, Diabetes, 38: 902–5.

Lopez-Otin, C., and J. S. Bond. 2008. ‘Proteases: multifunctional enzymes in life and disease’, J Biol Chem, 283: 30433–7.

MacLean, B., D. M. Tomazela, N. Shulman, M. Chambers, G. L. Finney, B. Frewen, R. Kern, D. L. Tabb, D. C. Liebler, and M. J. MacCoss. 2010. ’Skyline: an open source document editor for creating and analyzing targeted proteomics experiments’, Bioinformatics, 26: 966–8.

Martoglio, B., and B. Dobberstein. 1998. ‘Signal sequences: more than just greasy peptides’, Trends Cell Biol, 8: 410–5.

McAlister, G. C., D. P. Nusinow, M. P. Jedrychowski, M. Wuhr, E. L. Huttlin, B. K. Erickson, R. Rad, W. Haas, and S. P. Gygi. 2014. ‘MultiNotch MS3 enables accurate, sensitive, and multiplexed detection of differential expression across cancer cell line proteomes’, Anal Chem, 86: 7150–8.

Mentlein, R., B. Gallwitz, and W. E. Schmidt. 1993. ‘Dipeptidyl-peptidase IV hydrolyses gastric inhibitory polypeptide, glucagon-like peptide-1(7-36)amide, peptide histidine methionine and is responsible for their degradation in human serum’, Eur J Biochem, 214: 829–35.

Mulvihill, E. E., and D. J. Drucker. 2014. ‘Pharmacology, physiology, and mechanisms of action of dipeptidyl peptidase-4 inhibitors’, Endocr Rev, 35: 992–1019.

Okondo, M. C., S. D. Rao, C. Y. Taabazuing, A. J. Chui, S. E. Poplawski, D. C. Johnson, and D. A. Bachovchin. 2018. ‘Inhibition of Dpp8/9 Activates the Nlrp1b Inflammasome’, Cell Chem Biol, 25: 262–67 e5.

Ostapcuk, V., F. Mohn, S. H. Carl, A. Basters, D. Hess, V. Iesmantavicius, L. Lampersberger, M. Flemr, A. Pandey, N. H. Thoma, J. Betschinger, and M. Buhler. 2018. ‘Activity-dependent neuroprotective protein recruits HP1 and CHD4 to control lineage-specifying genes’, Nature, 557: 739–43.

Perez-Riverol, Y., J. Bai, C. Bandla, D. Garcia-Seisdedos, S. Hewapathirana, S. Kamatchinathan, D. J. Kundu, A. Prakash, A. Frericks-Zipper, M. Eisenacher, M. Walzer, S. Wang, A. Brazma, and J. A. Vizcaino. 2022. ‘The PRIDE database resources in 2022: a hub for mass spectrometry-based proteomics evidences’, Nucleic Acids Res, 50: D543–D52.

Petrella, L. N. 2014. ‘Natural variants of C. elegans demonstrate defects in both sperm function and oogenesis at elevated temperatures’, PLoS One, 9: e112377.

Schindelin, J., I. Arganda-Carreras, E. Frise, V. Kaynig, M. Longair, T. Pietzsch, S. Preibisch, C. Rueden, S. Saalfeld, B. Schmid, J. Y. Tinevez, D. J. White, V. Hartenstein, K. Eliceiri, P. Tomancak, and A. Cardona. 2012. ‘Fiji: an open-source platform for biological-image analysis’, Nat Methods, 9: 676–82.

Soneson, Charlotte, Vytautas Iesmantavicius, Daniel Hess, Michael Stadler, and Jan Seebacher. 2023. ‘einprot: flexible, easy-to-use, reproducible workflows for statistical analysis of quantitative proteomics data’, Journal of Open Source Software, 8: 5750.

Tagore, D. M., W. M. Nolte, J. M. Neveu, R. Rangel, L. Guzman-Rojas, R. Pasqualini, W. Arap, W. S. Lane, and A. Saghatelian. 2009. ‘Peptidase substrates via global peptide profiling’, Nat Chem Biol, 5: 23–5.

Taunton, J., C. A. Hassig, and S. L. Schreiber. 1996. ‘A mammalian histone deacetylase related to the yeast transcriptional regulator Rpd3p’, Science, 272: 408–11.

Tinoco, A. D., D. M. Tagore, and A. Saghatelian. 2010. ‘Expanding the dipeptidyl peptidase 4-regulated peptidome via an optimized peptidomics platform’, J Am Chem Soc, 132: 3819–30.

Tyanova, S., T. Temu, P. Sinitcyn, A. Carlson, M. Y. Hein, T. Geiger, M. Mann, and J. Cox. 2016. ‘The Perseus computational platform for comprehensive analysis of (prote)omics data’, Nat Methods, 13: 731–40.

Unhavaithaya, Y., T. H. Shin, N. Miliaras, J. Lee, T. Oyama, and C. C. Mello. 2002. ‘MEP-1 and a homolog of the NURD complex component Mi-2 act together to maintain germline-soma distinctions in C. elegans’, Cell, 111: 991–1002.

Wang, Y., F. Yang, M. A. Gritsenko, Y. Wang, T. Clauss, T. Liu, Y. Shen, M. E. Monroe, D. Lopez-Ferrer, T. Reno, R. J. Moore, R. L. Klemke, D. G. Camp, 2nd, and R. D. Smith. 2011. ’Reversed-phase chromatography with multiple fraction concatenation strategy for proteome profiling of human MCF10A cells’, Proteomics, 11: 2019–26.

Wilson, C. H., H. E. Zhang, M. D. Gorrell, and C. A. Abbott. 2016. ‘Dipeptidyl peptidase 9 substrates and their discovery: current progress and the application of mass spectrometry-based approaches’, Biol Chem, 397: 837–56.

Wingfield, P. T. 2017. ‘N-Terminal Methionine Processing’, Curr Protoc Protein Sci, 88: 6 14 1-6 14 3.

Xiao, Q., F. Zhang, B. A. Nacev, J. O. Liu, and D. Pei. 2010. ‘Protein N-terminal processing: substrate specificity of Escherichia coli and human methionine aminopeptidases’, Biochemistry, 49: 5588–99.

Yu, D. M., K. Ajami, M. G. Gall, J. Park, C. S. Lee, K. A. Evans, E. A. McLaughlin, M. R. Pitman, C. A. Abbott, G. W. McCaughan, and M. D. Gorrell. 2009. ‘The in vivo expression of dipeptidyl peptidases 8 and 9’, J Histochem Cytochem, 57: 1025–40.

Yu, D. M., T. W. Yao, S. Chowdhury, N. A. Nadvi, B. Osborne, W. B. Church, G. W. McCaughan, and M. D. Gorrell. 2010. ‘The dipeptidyl peptidase IV family in cancer and cell biology’, FEBS J, 277: 1126–44.

